# Active search signatures in a free-viewing task exploiting concurrent EEG and eye movements recordings

**DOI:** 10.1101/2022.06.27.497224

**Authors:** Damián Care, María da Fonseca, Matias J. Ison, Juan E. Kamienkowski

**Affiliations:** Laboratorio de Inteligencia Artificial Aplicada, Instituto de Ciencias de la Computación (Consejo Nacional de Investigaciones Científicas y Técnicas - Facultad de Ciencias Exactas y Naturales, Universidad de Buenos Aires), Argentina; Universidad Nacional de Río Negro, Argentina; School of Psychology, University of Nottingham, United Kingdom; Maestría de Explotación de Datos y Descubrimiento del Conocimiento, Facultad de Ciencias Exactas y Naturales, Universidad de Buenos Aires, Argentina; Departamento de Computación, Facultad de Ciencias Exactas y Naturales, Universidad de Buenos Aires, Argentina

**Keywords:** visual search, eye tracking, fixation-related potentials, face processing, deconvolutional analysis

## Abstract

Tasks we often perform in our everyday lives, such as reading or looking for a friend in the crowd, are seemingly straightforward but they actually require the orchestrated activity of several cognitive processes. Free-viewing visual search requires a plan to move our gaze on the different items, identifying them, and deciding on whether to continue with the search. Little is known about the electrophysiological signatures of these processes in free-viewing since there are technical challenges associated with eye movement artifacts. Here we aimed to study how category information, as well as ecologically-relevant variables such as the task performed, influence brain activity in a free-viewing paradigm. Participants were asked to observe/search from an array of faces and objects embedded in random noise. We concurrently recorded EEG and eye movements and applied a deconvolution analysis approach to estimate the contribution of the different elements embedded in the task. Consistent with classical fixed-gaze experiments and a handful of free-viewing studies, we found a robust categorical effect around 150 ms in occipital and occipitotemporal electrodes. We also report a task effect, more negative in posterior central electrodes in visual search compared to exploration, starting at around 80 ms. We also found significant effects of trial progression, and an interaction with the task effect. Overall, these results generalise the characterisation of early visual face processing to a wider range of experiments and show how a suitable analysis approach allows to discern among multiple neural contributions to the signal, preserving key attributes of real-world tasks.

## Introduction

Real-world free-viewing tasks require orchestrated sequences of events. When reading a sentence or exploring a visual scene, we move our eyes gazing on different elements of the scene creating a sequence of fixations alternated by saccade movements. The dynamic process that guide human saccadic scanpaths is known to be influenced by bottom-up factors such as the stimulus salience (Itti et al., 1998), but also by top-down factors such as the task (Castelhano et al., 2009), context (Torralba et al., 2006) and the semantic relations within the image (Underwood & Foulsham, 2006). In free-viewing tasks, these factors contribute to the ensemble of eye movements through the visual scene (Devillez et al., 2017; Devillez, Guerin-Dugue, et al., 2015).

One of the most informative stimuli we can find in real life is a human face. Indeed, a brief exposure to a human face is enough to gather crucial information such as its identity and emotion, and the EEG signal for faces show a consistent N170, an occipitotemporal component arising about 170 ms after stimulus presentation that is stronger for faces in comparison to objects (Joyce & Rossion, 2005; Rossion & Jacques, 2011).

On the other hand, studies investigating the brain processes underlying free-viewing are more scarce, since there are technical challenges associated with the eye movement artifacts in the EEG signal and the self-organised structure of the trial. However, through simultaneous EEG and eye-tracking (ET) recordings, it is possible to examine the neural processing underlying the visual information by extracting the fixation-related potentials (FRPs). Several studies involving free-viewing visual search tasks have shown components that distinguish between targets and distractors when looking for a special character (Kamienkowski et al., 2012; Hiebel et al., 2018) a face in a crowd (Kaunitz et al., 2014), or an object within a natural scene (Devillez, Guyader, et al., 2015). In this context of free-viewing, recent evidence indicates target detection is affected by refixation behaviour (Meghanathan et al., 2020), as well as the processing of semantic integration of the scene information (Coco et al., 2020). More recently, Auerbach-Asch et al. (2020) have included the dynamic of eye-movements in the context of free-viewing tasks, showing a similar topography of the N170 in comparison with a fixed gaze paradigm.

Most of these studies have manipulated an individual dimension in free-viewing. However, real-world free-viewing tasks require several concurrent processes to be operating in an orchestrated manner, rendering its analysis a challenge. Deconvolution models have extensively been applied in other fields such as fMRI (Dale & Buckner, 1997) and extended to EEG (Smith & Kutas, 2015). This framework has recently been applied to concurrent EEG and eye movements recordings using the Unfold toolbox (Dimigen & Ehinger, 2021; Ehinger & Dimigen, 2019) to study the impact of pre-saccadic preview over the post-saccadic face processing (Buonocore et al., 2020), as well as face selectiveness of neural activity in free-viewing (Auerbach-Asch et al., 2020).

In this study, we apply a deconvolution framework to decipher the contribution of different aspects of the task to free-viewing. We designed a paradigm for concurrent EEG and eye movements recordings, where faces and objects are embedded in a noisy background, and ask participants to perform two types of trials: *visual search* (VS) trials, in which they are instructed to search for a target, and *exploration* (EX) trials, in which participants observe the display with no explicit task. Our aim is to understand how the category (either a face or an object), the task (visual search or exploration), and the integration of information throughout the trial affect the neural responses.

## Materials and Methods

### Participants

Twenty-one adult participants performed the experiment. All participants were naïve to the objectives of the experiment, had normal or corrected to normal vision, and provided written informed consent, which was approved by the University of Nottingham School of Psychology Ethics Committee (ethics approval: F992). Data from five participants were excluded: two due to bad performance, two due to poor eye tracking quality, and one due to poor EEG quality even after filtering and applying the ICA procedure (see below). In total, we analysed data from sixteen participants with a median age of 23 years old (range = [19, 28] years old).

### Apparatus and data acquisition

Stimuli were presented on a BenQ XL2420Z monitor with a screen resolution of 1024 x 768 pixels and at a refresh rate of 75Hz. Participants were placed at a distance of approximately 60 cm from the monitor, and responses were introduced through a standard ‘qwerty’ keyboard. All stimuli were presented in MATLAB (MathWorks, 2000) using the psychophysics toolbox (Brainard, 1997).

EEG activity was recorded using a BioSemi Active-Two system at 1024 Hz with 64 electrode positions on an extended 10-20 montage. During data collection the Biosemi CMS electrode was used as reference. The CMS electrode was located between electrodes POz and PO3. As described by the manufacturer’s website, the location of the CMS does not influence the amplitude of the measuring electrodes (see manufacturer webpage for more details on the CMS/DRL system^1^. Four extra electrodes were placed on the right and left mastoids and the right and left earlobes. After data were recorded, the sampling rate was digitally downsampled to 512 Hz and the signal was re-referenced to the average of the 64 electrodes. Eye-movement tracking (ET) data were recorded using an EyeLink 1000 Plus system in monocular remote mode and a sampling rate of 500 Hz. A sticker with a bullseye (EyeLink target sticker) was placed on the forehead, between the eyebrows, as suggested by Eyelink developers. The eye-tracker system detects the sticker and uses it for head movement stabilisation. To integrate the ET and EEG data, the stimulus-generating computer sent shared messages to both the EyeLink and BioSemi systems at different temporal marks according to the experimental procedure.

### Experimental procedure

The experiment consisted of 9 blocks containing 24 trials each, making a total of 216 trials; the complete protocol lasted approximately 50 minutes. For all the experiments a drift correction was made at the beginning of every block, followed by a recalibration of the ET. Each block was composed of trials of the same type, either *visual search* (VS) or *exploration* (EX), (see Fig. 1). The main phase of each task (VS or EX), lasted 4.5 sec, regardless of whether the target had been found or not (VS). Six VS blocks and three EX blocks were run, mixed in a random order. We used a pre-experiment eye movement task “eye-map”, as in Ploch et al. (2012), to improve the artifact-correction procedure. This approach aims at getting data from intervals including saccades and blinks at times when the participant is not involved in the main task (therefore, minimising the risk of mixing artifactual and neural sources related to the task). Participants completed three eye-map blocks during the session. These were used in pre-analysis to overweight artifactual effects (Dimigen, 2020). In the eye-map task, there were five types of trials. In the first four, participants were asked to perform saccades between two horizontal or vertical points, separated by small or long distances. The dots remained present during the whole trial and participants were instructed to perform saccades at a slow pace, i.e. maintaining the fixation for some time before moving the eyes to the next location. The fifth condition was a single point in the centre of the screen where participants were asked to fixate and then perform blinks also at a slow pace. Event-marks were sent to both EEG and ET for offline synchronisation at the beginning and the end of each eye-map trial, as well as at the beginning of the eye-map block. Written instructions were displayed at the beginning of the experiment.

**Figure 1.**
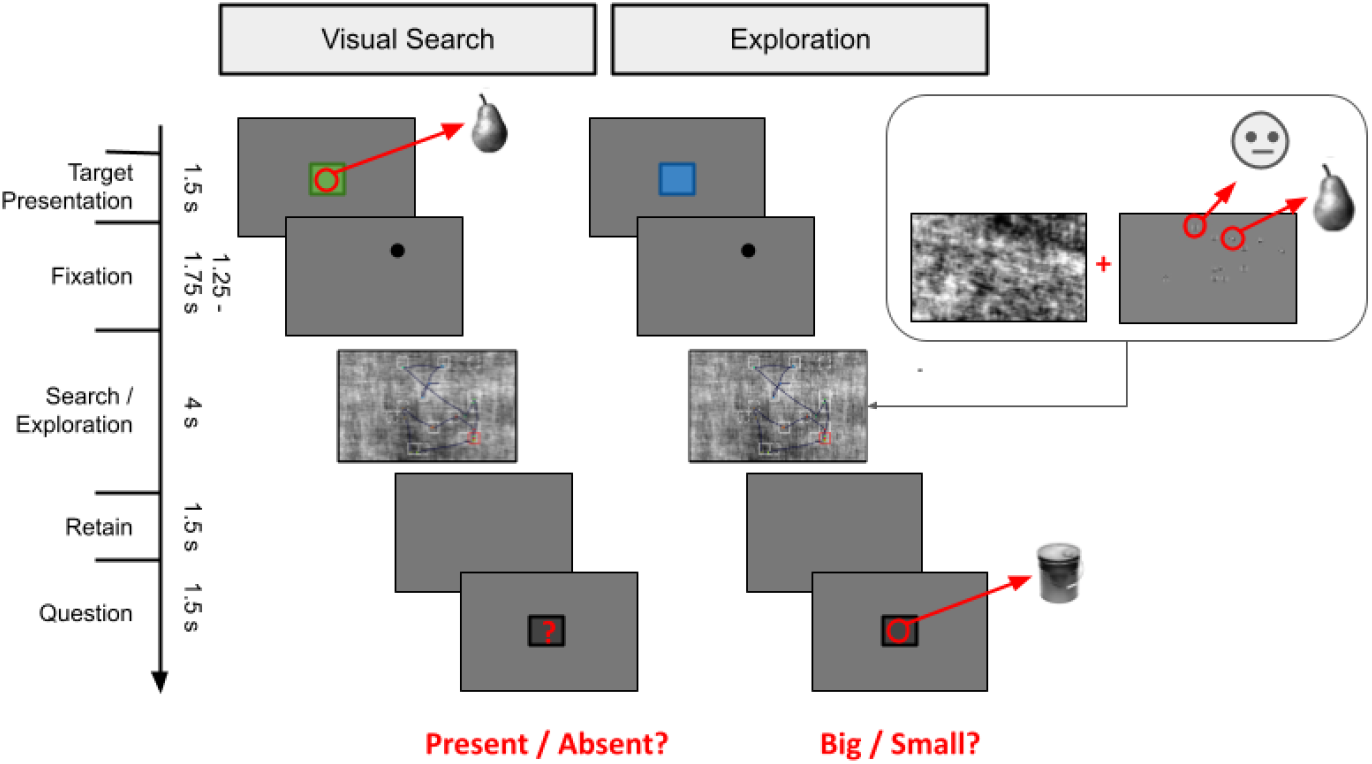
Experimental paradigm. Time progression of the trial for the Visual Search condition with target presentation and the Exploration condition with a final question unrelated to stimuli. In the search/exploration stage, the white squares indicate the position of the objects/faces, the red square indicates the position of the target, and the line is a scanpath. THIS FIGURE WAS MODIFIED FOR THE BIORXIV VERSION, IN THE ORIGINAL EXPERIMENT THE SMILEY FACE IS REPLACED BY A HUMAN FACE.

The VS task consisted of a search for a target image (face or object). Trials with each target category were inter-mixed in pseudo-random order within each search block. A picture of the target was displayed in the centre of the screen for 1.5 seconds. Immediately after, a fixation dot placed at a random position was exhibited on the screen for a lapse that lasted between 1.25 and 1.75 seconds. After the fixation phase, a crowd of 15 stimuli pictures (5 objects and 10 faces, see next section) immersed in a noisy background were exhibited for 4 seconds, and participants were instructed to search for the target image in the crowd. Participants were instructed to retain the information for a 1.5-second period in which a grey screen was displayed. Following the retain phase, a question appeared for 1.5 seconds asking the participants to press the left (right) arrow of the keyboard to answer if the target image was present (absent) in the crowd. The target was present in the crowd of items half of the time. A correct answer to this question marked the trial as correct. Target presentation, fixation, retain and question phases were presented in front of a uniform grey background image.

The EX task involved a procedure similar to the VS task. First, an empty box was displayed in the centre of the screen for 1.5 seconds. Immediately after, the same fixation, exploration and retain phases of VS task were presented, with the only instruction for free exploration of items in the crowd. At the end, in the question phase, either a face or an object was displayed in the centre of the screen with a question underneath asking, for example, whether the object is big (left arrow key) or small (right arrow key). This item was always absent in the exploration phase. Before the experiment, the participants classified several pictures of objects as big or small compared with a football. Trials were tagged as correct in accordance with the reply to this question.

### Stimuli

A set of 369 face grey-scale images were obtained from the Aberdeen face database^2^, Karolinska directed emotional faces (Lundqvist *et al*., 1998)^3^, Yale face database^4^, the AT&T face database^5^, the Face-Place database^6^, and the Aberdeen, the Iranian, the Pain and the UTrecht databases from the Psychological Image Collection at Stirling site (PICS)^7^. Face stimuli were carefully selected to have neutral expressions. Another set of 821 object grey-scale images were selected from the Unique Objects database (Brady et al., 2008)^8^. All stimuli were framed in 250 x 250 pixels boxes and made isoluminant with the background (Supp. Fig. 2).

### EEG and eye movements pre-processing

EEG data were imported, pre-processed and analysed with MATLAB using the EEGLAB toolbox (Delorme & Makeig, 2004) and custom-made scripts. Firstly, EEG raw data was filtered in the range 0.1-100 Hz and a notch filter was applied at power grid line frequency. Noisy channels with abnormal spectra were interpolated as long as there were less than five noisy spatially contiguous channels. We excluded one participant who didn’t meet this criterion Then, eye tracking and EEG data were synchronised with the EYE-EEG toolbox (Dimigen et al., 2011) using the synchronisation marks. Gaze positions landing outside screen range and blinks were marked as bad data. Eye movements were recalculated using an offline algorithm (Engbert & Mergenthaler, 2006).

To remove eye movements artifiacts we used an approach based on Independent Component Analysis (ICA), as previously done in experiments involving concurrent EEG and eye movements recordings (Buonocore et al., 2020; Dimigen, 2020; Kamienkowski et al., 2018; Keren et al., 2010; Ossandón et al., 2010; and others). In such studies, the EEG signal is aligned to the onset of fixations or the onset of saccades leading to fixation-related potentials (fRPs) or saccade-related potentials (sRPs). This allows researchers to investigate the brain activity elicited on each fixation. We have previously used this approach (see Kaunitz et al., 2014) to show a high similarity between fRPs recorded in an eye movement visual search task and ERPs recorded in a fixed-gaze experiment with similar stimuli. One major obstacle, however, arises from the fact that the neural signals in the EEG are contaminated by non-cerebral electrical sources. As elegantly described by Keren et al., (2010), the three main ocular artifacts (eyelid movement, corneo-retinal dipole (CRD) rotation, and an ocular myoelectric signal called saccadic spike potential or SP) differ in their spatial, temporal, and spectral signatures. Typically, the amplitude of these artifacts is higher than the neural signal of interest (occuring during fixations) and their removal is desirable (Dimigen, 2020). There are several methods that have been proposed to detect ocular artifacts, including regression approaches using the EOG electrodes (Croft & Barry, 2000; Parra et al., 2005) and ICA. Prior to ICA training, as suggested by the OPTICAT procedure (Dimigen, 2020), the perisaccadic samples were overweighted by expanding the original data to include a short [-30 30] ms interval around the saccades. This step, which is performed before ICA training, has been shown to improve the quality of ICs representing the myogenic saccadic spike potentials, therefore resulting in improved artifact correction (Keren et al., 2010). Afterwards, ICA was calculated with Infomax algorithm (Bell & Sejnowski, 1995) and principal component analysis (PCA) was used prior to training in those cases in which data included interpolated channels. Next, we applied a variance criteria (Plöchl et al., 2012) to recognise components related to ocular artifacts, a manual selection of ICA projections was also performed to exclude components in which OPTICAT failed to automatically recognise ocular movement sources. This manual step was only needed in less than 6.6% (SD: 2.5%, max: 10.9%) of the total number of components. Lastly, the 0.2 high-pass EEG data was reconstructed using the weights of the surviving ICA components calculated on the 2 Hz high-pass filtered data.

For each analysis, in the case of micro-fixations, that is, successive fixations to the same item, we selected the first micro-fixation in the array of successive micro-fixations that exceeded 100 ms duration as the event-related fixation. In the case of non-successive fixations to the same item, we only kept the first and the second one. We discarded the first fixation of each trial, and the trials with less than three fixations. In order to compare the neural data associated with the fixation behaviour during active search and free exploration, we kept fixations within the correct trials and fixation duration within the range [100, 1000] ms. We defined *active search* (AS) as the set of fixations to non-target items belonging to the VS task with the target present, previous to finding the target (first fixation to target longer than 200 ms).

### Single-participant (first level) statistical analysis

For FRP construction, we ran mass univariate regressions on the continuous data with the Unfold toolbox (Ehinger & Dimigen, 2019). This was done for each participant separately. We defined the time interval of interest around each fixation onset as the time from 200 ms before to 400 ms after the fixation event; this interval was used to build the design matrix and determine the time limits of the analysis. Fixations marked as bad data were discarded. We used treatment coding for categorical effects (e.g. ‘task’, ‘category’) which consists of dummy-coding (with ones and zeros) on all but one of the levels of the factor (reference level with zero value) (Smith & Kutas, 2015). Since the intercept term is the estimation of the FRP when all variables are zero, the interpretation of the parameters depends on the base condition. For instance, deviance at 200 ms in the time course of an estimated parameter would mean an effect at that very latency resulting from the difference between the FRP associated with the considered parameter and the FRP associated with the reference level. For a continuous variable, we interpret the values of the estimated parameters as the change in the predicted FRP per unit of the continuous variable. Furthermore, interaction parameters give information about non-additive interactions between factors (See Supplementary Methods for details on the interpretation).

The effect associated with the *category* (non-target objects: NTO as a reference level, non-target faces: NTF) on the FRPs was analysed following this model:

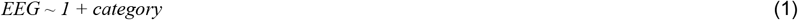

Where we adopted ‘symbolic’ notation to specify the model (see SI for the specification of our models in a different notation). To analyse the effect related to the *task*, we introduced a dummy variable indicating whether the condition is AS or EX (as reference level). The corresponding formula was:

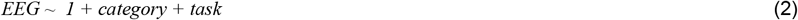

Then, we incorporated the fixation rank (centred to the mean, and standardised per trial) as a continuous variable named *trial progression score* (TPS):

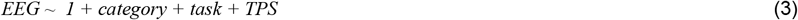

Lastly, we introduced the interaction between the *task* and the *TPS* in order to capture the trial progression score trend in each condition

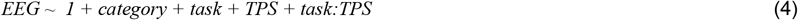

This analysis was done for each participant and electrode separately. We used the equations 1-4, with the variables of interest, and 74 delays, from -200ms to 400ms, to build the expanded design matrices. When expanding the model, we only kept significant variables. We regressed continuous data with the expanded design matrix and obtained one beta value (or estimate) for each variable and delay (Ehinger & Dimigen, 2019). The resulting beta values are called regression ERPs (rERPs) (Smith & Kutas, 2015), or rFRPs in our case, or temporal response functions (TRF) (Coppola, 1979; Crosse et al., 2016; Dandekar et al., 2012; Lalor et al., 2009). These were treated similarly to FRPs. We applied a baseline correction to the obtained responses with a baseline period taken from 200 ms before each fixation and ending at fixation onset. After that, we performed second-level analysis of these TRFs obtained for each participant and variable to assess the significance of the responses, correcting for multiple comparisons.

### Group (second-level) statistical analysis

Second-level statistical analysis was carried out using the threshold-free cluster enhancement method (TFCE) (Mensen & Khatami, 2013; Smith & Nichols, 2009). The TFCE is a data-driven approach that uses permutation-based statistics for the analysis of EEG data. As other cluster correction approaches (Bianchi et al., 2019; Maris & Oostenveld, 2007; Mensen & Khatami, 2013; Pernet et al., 2015), it takes into account the dependence between neighbouring EEG samples, both in time and space, and results in a much less conservative approach than t-testing with Bonferroni correction. It was first introduced as an improvement to cluster-based permutation tests (where an arbitrary threshold value affects the sensitivity of clusters of different sizes and intensities (Holmes et al., 1996; Smith & Nichols, 2009), first applied to fMRI analysis (Holmes et al., 1996; Smith & Nichols, 2009) and later extended to EEG (Mensen and Khatami 2013). We used Mensen’s MATLAB implementation of TFCE (Mensen & Khatami, 2013)^9^.

The procedure was conducted after the first-level analysis, where TRFs are obtained for each variable and participant. These TRFs, which are composed of the beta values or slopes for each electrode and time, were then compared against zero (null hypothesis of no dependence between the EEG and the variable of interest in each electrode and sample). As we performed one-sample tests, we randomly changed the sign of the slopes (beta values). This is the same approach used by Coco et al. (2020) among others, and suggested by the FieldTrip toolbox. This procedure is repeated N=2000 times for each electrode and time-sample to build a set of 2D matrices under the null hypothesis. Only *p*-values under 0.05 are shown for visualisation of the TFCE results.

## Results

In this study, we developed an experiment involving data from concurrent EEG and eye movements recordings. Stimuli were composed by a set of images of objects and faces embedded in a noisy background. During the experiment, participants performed two types of tasks split into different trials: *visual search* (VS) trials, in which they searched for a target, and *exploration* (EX) trials, where they just explored stimuli presented on the screen. By keeping only the correct target-present trials of the VS trials and the fixations previous to the target, we studied the *active search* (AS) behaviour and compared it with the behaviour during EX trials. The overall task performance was 78% in VS (present/absent target question after the search screen disappears) and 82% in EX trials (big/small stimulus question after the exploration screen disappears).

### Eye-movements behaviour

The eye movements showed the typical characteristics of free-viewing experiments (see Fig. 2). Saccade amplitude and main sequence of saccades (Bahill et al., 1975; Harris & Wolpert, 2006) followed a typical distribution (Fig. 2B,E) associated with the non-linearity in the saccadic system (Lebedev et al., 1996) considered to be present in both types of tasks (Otero-Millan et al.,2008). Saccade direction in VS condition showed a horizontal preference (Fig. 2C) which was slightly expanded in the EX condition (Fig. 2F). Fixation duration distribution was also typical, with a peak at around 170 ms, and a mean duration of (195 ± 110) ms (Fig. 2G). These results were consistent with previous findings in visual search and exploration (Otero-Millan et al., 2008). Fixation duration to targets was longer than to non-target stimuli during VS condition (Fig. 2H). Finally, the mean number of fixations per trial was 8 with a standard deviation of 5.

**Figure 2.**
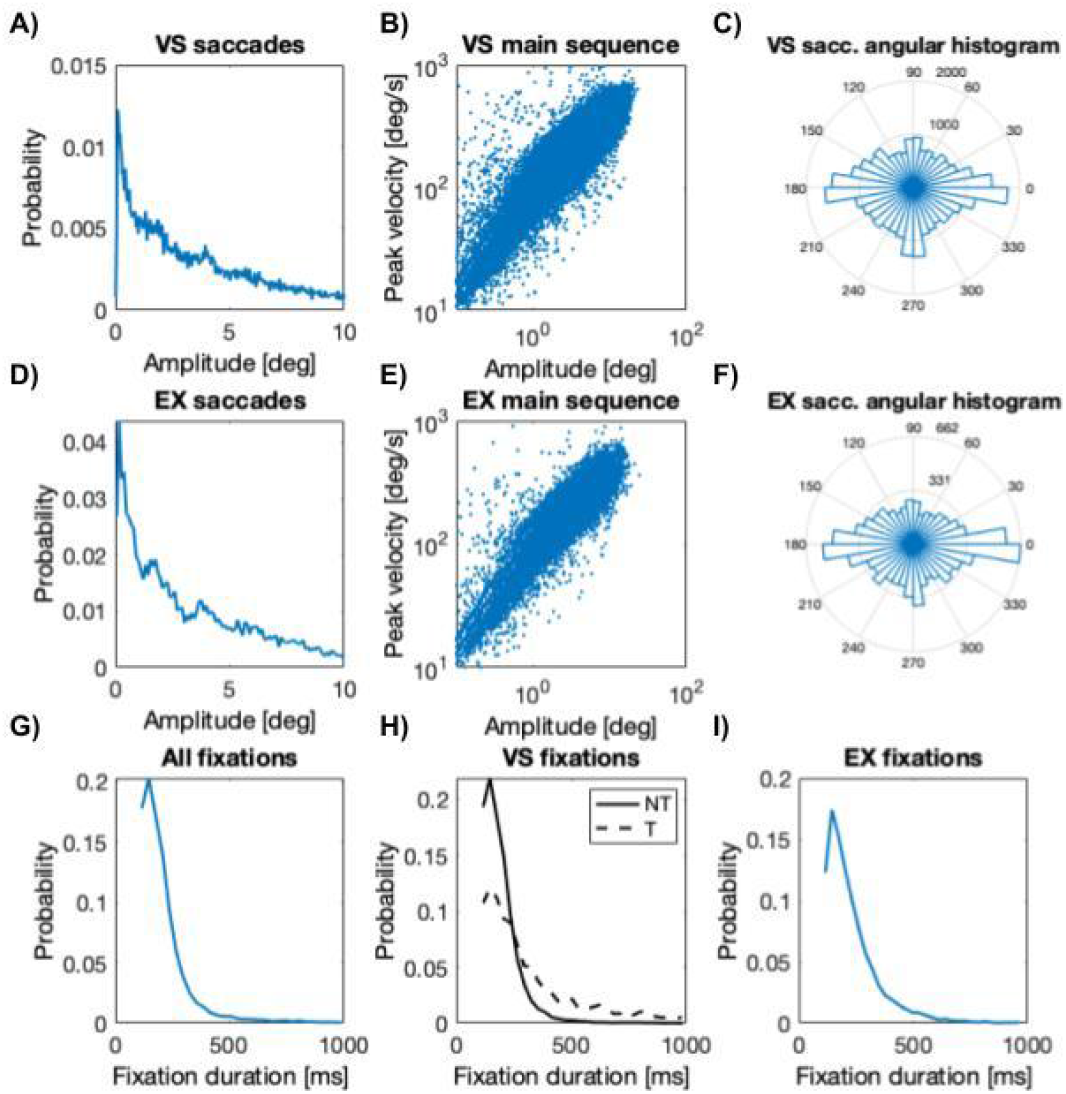
Eye movement behaviour for all participants. **A. D**. Saccade amplitude distribution in degrees of visual angle, **B. E**. Main sequence of saccades, **C. F**. Saccade angular distributions, **G**. fixation duration distribution for all saccades from all the trials included in the analysis (N=8654), **H**. fixation duration distribution for all saccades from the visual search (VS) trials, separated by identity: target (T, bold line) and non-target (NT, dashed line), and **I**. fixation duration distribution for all saccades from the exploration (EX) trials.

Sometimes, when a participant fixates a stimulus, microsaccades occur within the fixation (i.e. a multiple fixation). We established a criterion to determine which of those multiple fixations on the same stimulus would be marked as a fixation event to extract the EEG responses. Here, we aim to get the best representation of the response to the stimulus that captured the participant’s attention. To that end, we only considered EEG responses to the first fixation within those multiple fixations whose duration was greater than 100 ms. For instance, if the first fixation lasted only 40 ms and the second lasted 150 ms, then we considered the second fixation as the one that would identify the response of interest.

### Non-Target Objects and Non-Target Faces

Firstly, we explored differences in the brain response to the category of the fixation content, in particular, we considered fixations to non-target objects and compared them with fixations to non-target faces. Because there could be an overlap between the evoked response of consecutive fixations, we applied a regression-based analysis to disentangle the overlapping activity (Ehinger & Dimigen, 2019). We estimated the response to the stimulus category, i.e. the difference between FRPs to faces and objects (Eq. 1). Fixations from both tasks, AS and EX, were included in these analyses.

The estimated response to fixations to non-target objects (intercept term) had a well-defined P1 component at 100 ms, there was a local minimum between 120 ms and 200 ms, and a second smaller positive component at around 230 ms (Fig. 3C). These activations were present in all occipital and occipitotemporal electrodes (Fig. 3B). The frontal components exposed an inverted polarity (Fig. 3B). On the other hand, the response to category exhibited an early effect from 130 ms to 300 ms. This effect, associated with the variable that encodes the category of the stimuli (face or object; Fig. 3C), is compatible with an N170 component. Similar results were obtained including post-target fixations (see Supp. Fig. 3).

**Figure 3.**
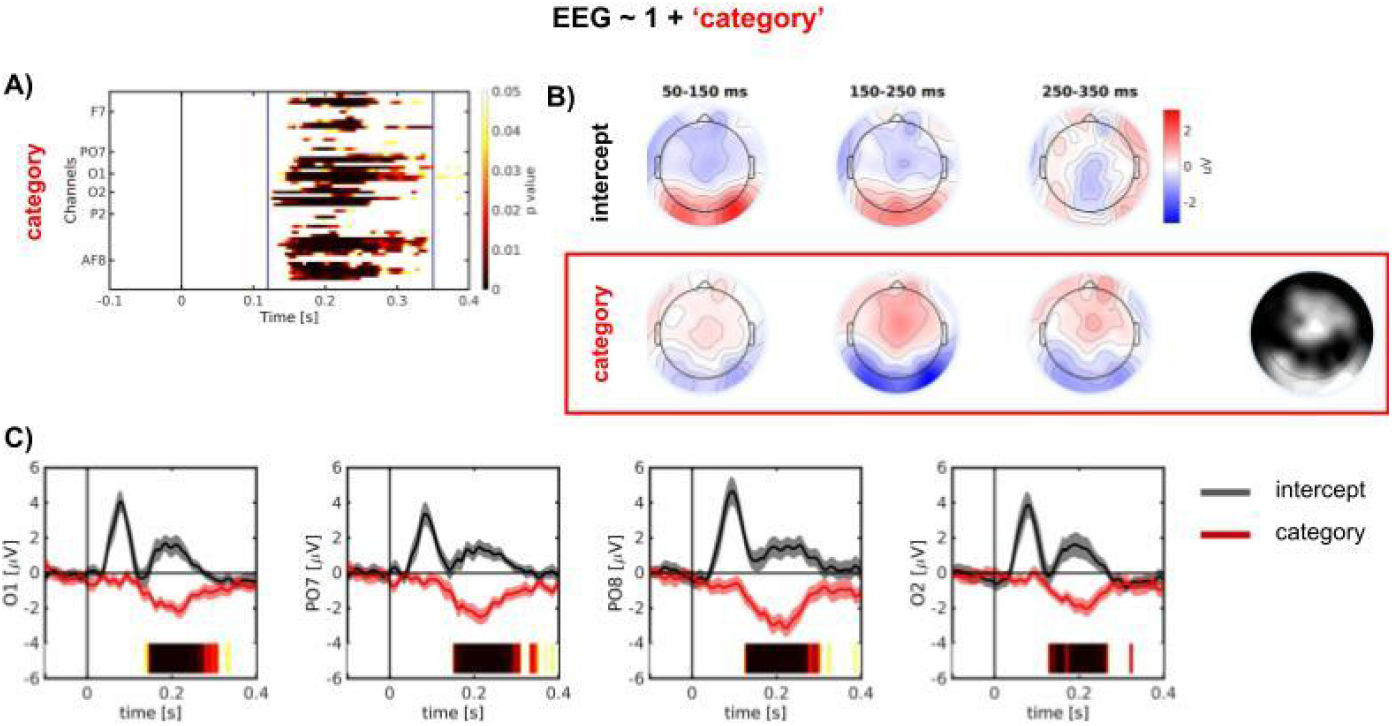
Time-course of category effect on FRPs for active search and exploration tasks (model Eq. 1) **A**. Significant samples obtained with the TFCE. Vertical lines represent the start and end of the significant cluster (with p<0.01). **B**. Scalp topography for different stages of the intercept and the category effect (beta values). The right scalp topography shows the proportion of significant samples per channel during the interval of the significant cluster (vertical bars in A). The colour scale represents activation from 0 (black) to the whole interval (white). **C**. Mean and s.e.m. across participants of the intercept (black) and the category (red) effect for selected channels [O1, PO8, PO7, O2]. The colour bar below each panel represents the significant intervals after TFCE second-order statistics applied to the face-object (category) effect. The intensity of the bar represents the significance level (p-value), only for the p-values < 0.05, the scale is the same as in panel A. The shades represent the standard error of the mean across participants. Number of fixations = 5127. Number of participants = 16.

We used the TFCE method to assess the statistical significance of FRPs associated with the stimulus’ category (Fig. 3A). We can see significant samples of this effect in central and occipital regions of the scalp that extend consistently from 120 to 350 ms (Fig. 3A,C).

### Early potentials evoked by task (between trials effect)

To explore how task changes influence FRPs, we included a binary variable representing the task (AS or EX) in the model (Eq. 2), and estimated the time course of the response. The task effect on the FRPs showed slightly earlier activation (green curve in Fig. 4, panel D) and a different topography (Fig. 4, panel C) compared to the content (face/object) effect. Scalp topographies showed that the task effect (VS - EX contrast) presented a negative occipital and a positive frontal component. Interestingly, a frontal activation was also present in the task effect, for which the TFCE yielded significant intervals from 80 ms to 250 ms (Fig. 4B,D). Regarding the category effect distribution, it remained essentially unchanged (Fig. 4C), in comparison with the simplest model (Fig. 3). The task effect showed a posterior central scalp distribution starting around 80 ms, while the category effect showed a bilateral negativity that started later. These differences can point towards a P1/lambda component modulation by the task, and an N170 component associated with faces. We also evaluated a model that includes an interaction term between Category and Task. As this interaction did not show any significant effects (see Supp. Fig. 4), this term was not included in larger models.

**Figure 4.**
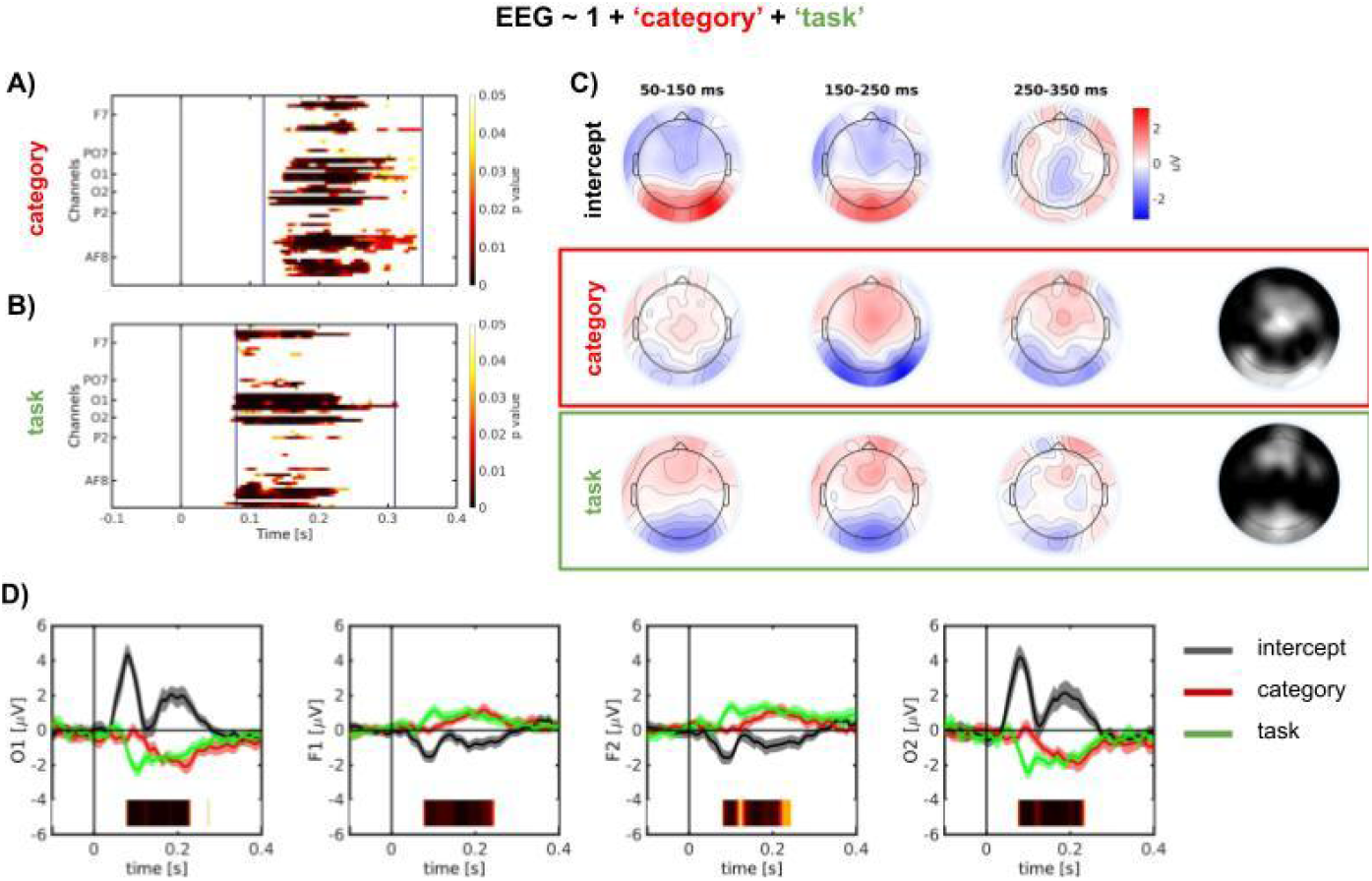
Time-course of task effect on FRPs for active search and exploration tasks (model Eq. 2). **A.B**. Significant samples obtained with the TFCE test for the category (**A**) and the task (**B**) effects. Vertical lines represent the start and end of the significant cluster (with p<0.01). **C**. Scalp topography for different stages of the intercept and the category effect (beta values). The right scalp topography shows the proportion of significant samples per channel during the interval of the significant cluster (vertical bars in A-B). The colour scale represents activation from 0 (black) to the whole interval (white). **D**. Mean and s.e.m. across participants of the intercept (black), the category (red) and the task (green) effect for selected channels [O1, F1, F2, O2]. The colour bar below each panel represents the significant intervals after TFCE second-order statistics applied to the AS-EX (task) effect. The intensity of the bar represents the significance level (p-value), only for the p-values < 0.05, the scale is the same as in panel A. The shades represent the standard error of the mean across participants. Number of fixations = 5127. Number of participants = 16.

### Global effects of the task and trial progression on the FRPs

Lastly, we included in the analysis the trial progression score (TPS) as a continuous variable through the standardised order of each fixation in the trials. To explore how these effects interacted with each other, we fitted a model that also included the interaction between the task and the trial progression score variable (Eq. 4). The present regression-based analysis showed the time course of the effects that varied between tasks and the TPS effect that varied within each trial, for a given task.

The FRPs of the content and task effects remained virtually unchanged. In the case of the TPS effect, FRP rose up to an almost constant value at about 170 ms for occipital channels, while frontal channels exhibited an opposite behaviour with a smoothly varying pattern around 200 ms (Fig. 5F). Second level statistics results showed a considerable occipital and frontal area of significance for these effects (Fig. 5C) starting at 100ms. The scalp topographies for the TPS effect exhibited a spatial distribution that largely remained the same until the end of the fixation period considered in the analysis (Fig. 5E). Similar results were obtained when fitting data with a model without the interaction term (Eq. 3; see Supp. Fig. 5).

**Figure 5.**
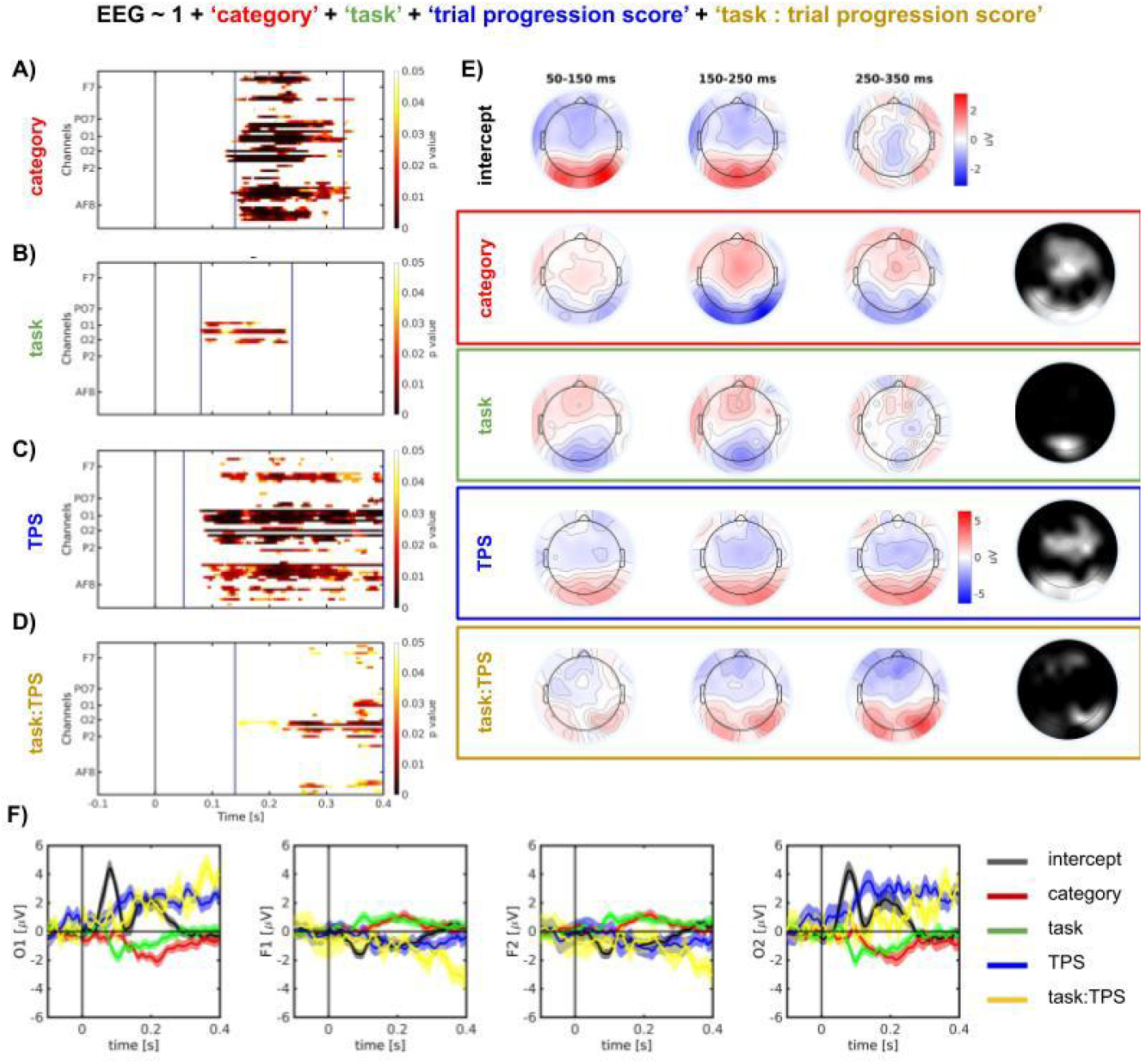
Time-course of trial progression score (TPS) and the interaction with the task effect on FRPs for active search and exploration tasks (model Eq. 4). **A.B.C.D**. Significant samples obtained with the TFCE test for the category (**A**), the task (**B**), the TPS effect (**C**), the interaction between TPS and the task (**D**). Vertical lines represent the start and end of the significant cluster (with p<0.01). **E**. Scalp topography for different stages of the intercept and the category effect (beta values). The right scalp topography shows the proportion of significant samples per channel during the interval of the significant cluster (vertical bars in A-D). The colour scale represents activation from 0 (black) to the whole interval (white). **F**. Mean and s.e.m. across participants of the task (green), the TPS (blue) and the interaction between the TPS and the task (yellow) effects for selected channels [O1, F1, F2, O2]. The shades represent the standard error of the mean across participants. Number of fixations = 5127. Number of participants = 16.

The interaction effect between the TPS and the task exposed a later activation that followed the tendency of the TPS effect, with positive occipital and negative frontal activation. The second-statistical level results for the responses to the interaction supported the idea that TPS was more influenced by the active search of a target (AS) in comparison with the free exploration of items (EX) (Fig. 5D).

Interestingly, the intercept presented a similar form through the different models, meaning that the estimate of the FRP is largely unchanged when additional factors are taken into account, such as task and or stimulus category.

The current framework allows to separate the contributions from successive fixations disentangling overlapping components. Recently, some prominent studies have used a similar approach to also separate the contributions of the saccades (Coco et al., 2020; Ossandon et al., 2020; Dimigen & Ehinger, 2021; Gert et al., 2022). We expanded the full model to also include an intercept and the saccade amplitude modelled with a fifth-order spline associated with the saccade onset. A direct comparison between the two models obtained by the TFCE test showed that both the significant electrodes in which each of the variables showed significant effects and the times in which this happened were largely preserved (Supp. Fig. 6).

Up to this point, we focused on active search (AS) defined as fixations before the target was found. In the target-present condition, this assures that the observer is indeed looking for the target. In the target-absent (TA) condition, the observer is also looking for the target for a potentially larger number of fixations. However, there could be a point in the trial in which the participant gives up the search and decides that the target is not present. To explore the active search intervals in TA trials, we defined fixation rank as the fixation position in the sequence (scanpath) and implemented a model including the first *N_threshold* fixations (i.e. *fixation rank < N_threshold*) in the TA condition during visual search (VS-TA), and the exploration (EX) trials (Fig. 6A) using the same variables of eq. 4 and Fig. 5.

**Figure 6.**
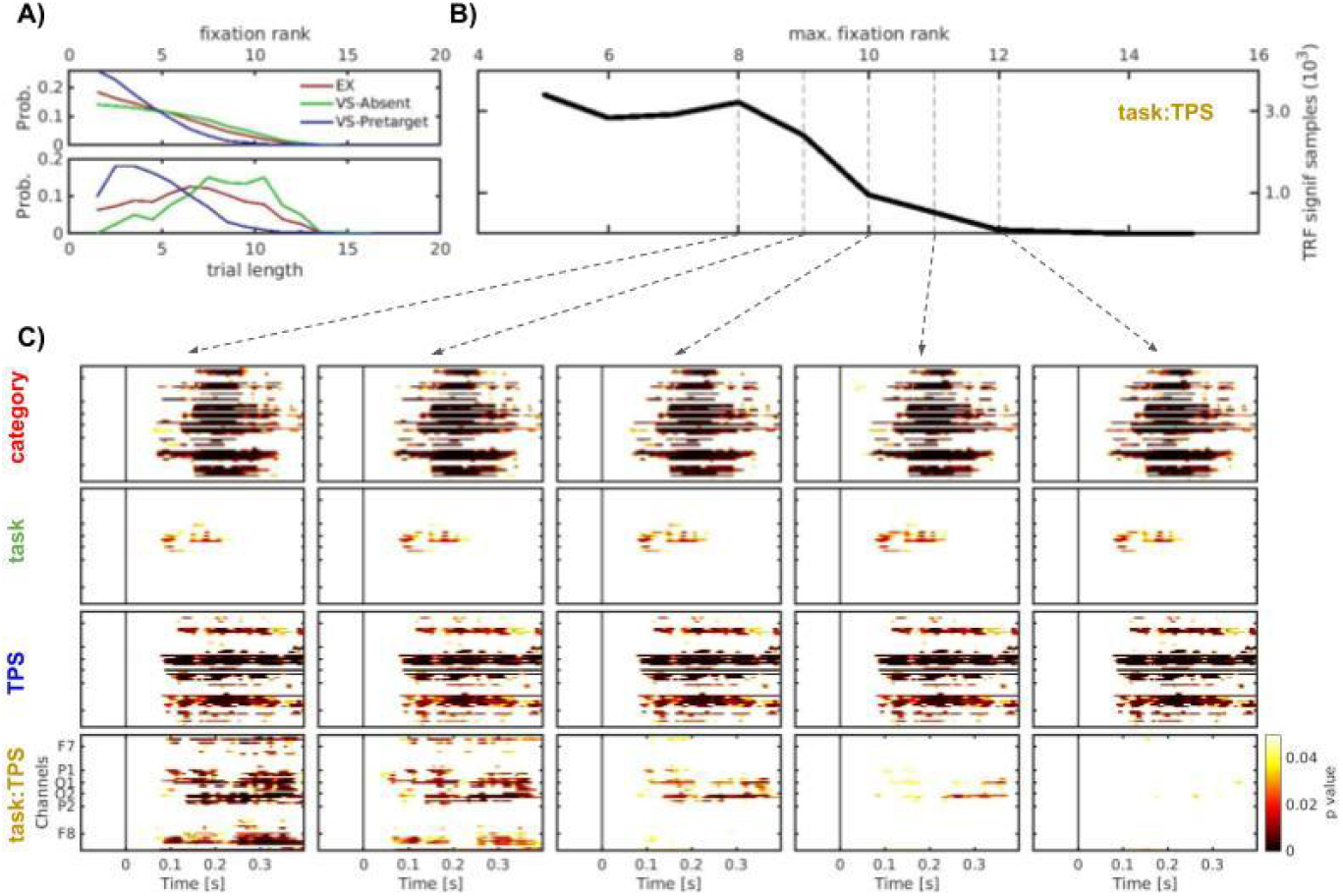
Target absent trials. Interaction between task and TPS. **A)** *Top:* Distribution of the fixation rank, i.e. the fixation position within the scanpath, during Exploration (EX; N=3292, median=4, max=13), VS-Target absent (VS-TA; N=3292, median=4, max=13), and fixations before the first fixation to the target (VS-Presentarget during AS; N=2333, median=4, max=14). *Bottom:* Distribution of the trial length (in fixations) for EX (N=626, median=7, max=13), VS-TA (N = 663, median = 9, max = 16) trials, and VS-Pretarget during AS (N = 630, median = 5, max = 14). In all cases only fixations to distractors were considered. **B)** Number of samples with significant task:TFS interaction, i.e. summarising the last row of panel C, as a function of the maximum fixation rank included in the analysis. **C)** Model significance after TFCE for each variable for the EX and VS-TA conditions, considering fixation ranks smaller than 8, 9, 10, 11, and 12. The general criteria for including fixations was similar to previous analyses: 1. only correct trials, 2. no refixations, 3. fixations with durations between 100 and 1000ms, and 4. excluding the first fixation of the trial. The models have the same variables as the full-model in Figure 5.

We studied VS-TA trials with 5 to 15 fixations, which allowed us to keep enough fixations for comparison, parametrically varying the fixation position within the trial (see Fig. 6A). Interestingly, the TRFs to all variables remained the same as in Fig. 5, with additional non-significant effects associated with the interaction between Rank and Task only when 12 or more fixations were considered (Fig. 6C). In order to summarise this effect, we counted the number of significant samples (time-by-channel) in the interaction between Rank and Task as a function of the *N_threshold* considered (Fig. 6B). This showed a clear transition from a significant effect (when the participant was expecting to find the target) to a non-significant effect (when, arguably, participants had given up the search).

These results are in line with our assumption that, at some point, the observer gave up the search and started wandering, similar to exploration. This is a very relevant observation to the dynamics of visual search, and opens the door to future studies, complementing neurophysiological measures with computational models able to extract hidden variables from behaviour, to determine the inflection point in which the observer quits the search (Moran et al., 2013).

## Discussion

In the present study, we aimed to characterise how, under free-viewing conditions, visual information processing depends on the content of the stimuli being fixated, the task being performed and its progression. We used a mass univariate deconvolution modelling approach to reduce the effects of overlapping brain activity in successive fixations, such as preparation and execution of eye movements and the visual stimulation itself. We modelled the resulting fixation-locked potentials with linear regressions by sequentially adding different variables (category of the fixated stimulus, task, and a trial progression score). Cluster-based permutation tests were used to control for multiple comparisons.

We observed a significant category (faces vs. objects) effect that resembles the well-known EEG N170 component seen in fixed-gazed experiments (Joyce & Rossion, 2005; Rossion & Jacques, 2011), fast periodic face stimulations (Liu-Shuang et al., 2014), and, more recently, in an experiment allowing eye movements in controlled displays (Auerbach-Asch et al., 2020). We started with a simple model that only included the category effect. Interestingly, this effect remained almost unchanged (both in latency and topography) when other variables were included in the model. This suggests an effect that is independent of the task and the progression of the trial and reflects the processing of the foveated item. Moreover, the short latency of the category effect (with a significant effect starting at 120-130 ms from fixation onset), is comparable to the one reported in rapid categorization tasks, with significant differences in the EEG response between categories at around 150 ms (Thorpe & Fabre-Thorpe, 2001; VanRullen & Thorpe, 2001). This was linked with saccadic response times in behavioural tasks (Kirchner & Thorpe, 2006) and further supported by intracranial recordings (Kirchner et al., 2009).

Then, we explored the effects of the task and the trial progression in the model, like variables that could affect FRPs throughout saccades, i.e. global variables. Firstly, we focused on the differences between active search (AS) and exploration (EX). The effects of the task in the FRPs have been previously explored in the context of modulating task demands on a guided-search task (Ries et al., 2016) and comparing free-viewing visual-search/memorization task versus the recall display (Seidkhani et al., 2017). Ries and collaborators (2016) asked participants to perform a guided-search task, moving their eyes towards the item marked with a red circle until they found the target. At the same time, they were asked to either follow or ignore the auditory information from a 0, 1, or 2-back task. Thus, the concurrent task modulated the resources available for the guided-search task. They observed that higher auditory task demands increased the latency and decreased the amplitude of the P3 in response to the target during the guided-search, while the lambda response showed a decrease in amplitude only in the high task demands condition (Ries et al., 2016). In our case, we manipulated the task itself instead of introducing a concurrent task and exploited the capabilities of the linear models to separate the effect of the early visual responses from the task. We observed that the task showed a significant effect that started early in the fixation (before 100ms) and spread to almost 250ms, while the specific responses to the category remained unaffected. This interval overlaps with the one observed by Seidkhani and collaborators (2017) while comparing the networks that emerge after the fixation onset between a search/encode phase and a recall phase in a change detection task that included free eye movements (Seidkhani et al., 2017).

The task manipulation could also affect the FRPs through an increase in temporal attention or a higher attentional engagement in active search (Corbetta et al., 1998; Correa et al., 2006; Kamienkowski et al., 2018; Melcher & Colby, 2008). For instance, targets appearing at attended moments in highly demanding perceptual processing tasks have an effect both on the P1 and the N2 components (Correa et al., 2006). However, these previous studies could not disentangle whether this was a modulation of the same component or a new effect, as we showed here by simultaneously analysing both effects. In one study, Carmel and Bentin (2002) approached this question by analysing the responses to Cars and Faces in different tasks (Carmel & Bentin, 2002). They found that the response to Faces (N170), remained the same across tasks, while the negative deflection in response to Cars at the same latency changed when cars were relevant to the task. They suggested that the visual mechanism involved in face processing is not influenced by task, while the processing of other stimuli, as part of a more general visual mechanism, is sensitive to strategic manipulations in attention. However, in our task, we didn’t find evidence of a relationship between task and category. The results presented here, with significant early task effects, might be associated with higher attentional engagement, where earlier responses are expected on higher perceptual load tasks in accordance with perceptual load theories (Lavie et al., 2014). Indeed, previous studies have reported that early task effects are more prominent in highly demanding tasks such as active search. We previously reported early target detection in tasks involving prolonged fixations when participants were asked to detect a subtle change in a synthetic stimulus (Kamienkowski et al., 2012) or when naturally looking for a face in a crowded scene -i.e. producing fixations in the order of 200-250ms (Kamienkowski et al., 2018), but not when they were instructed to prioritise accuracy in each fixation over time, producing longer fixations to avoid missing the target (Kaunitz et al., 2014). This is also consistent with studies reporting that target identification can be detected from the EEG signal prior to fixation onset (Dias et al., 2013; Stankov et al., 2021).

We have previously explored the effect of task progression in FRPs, which exhibited signatures related to the accumulation of evidence during the task (Kamienkowski et al., 2018). Here, the task progression showed a massive effect that started around the first 50ms and continued after 400 ms. Before 150 ms this effect reflected a local increase (with respect to the reference level) only in the occipital region, while from 150 ms it also exhibited a decrease in fronto-central electrodes. As mentioned before, this effect did not interact with the categorical response. Still, fixations in active search presented an even stronger pattern as shown by the interaction, providing evidence for the task demand when participants actively search for a target. Indeed, the steady decrease in central-frontal regions following the progression of the search was similar to the significant decrease in the baseline amplitude found in previous visual search free-viewing experiments (Kamienkowski et al., 2018), and to gradual changes in the baseline observed in other tasks in which participants were required to accumulate some evidence in order to achieve a decision, both in humans (de Lange et al., 2010; Pinheiro-Chagas et al., 2019) and monkeys (Yang & Shadlen, 2007).

The changes in brain activity as a function of fixation rank could also be interpreted as updates on the predictions on the potential target location (linked to the eye movement scanpaths) or on whether the target is present or not (linked to the decision). These processes are related to the integration of previous evidence that lead to the decision of when/where to move the eyes next or when to stop the search (and decide if the target is absent).

The stronger changes along the task in active search relative to exploration could also be due to increased working memory, general cognitive demands, or frustration. Indeed, as shown by Summerfield and others (Summerfield et al., 2008; Summerfield & Egner, 2009), neural adaptation is strongly connected to top-down perceptual expectations and selective attention. Previous works showed how visual working memory capacity is strongly related to performance in both visual search without eye movements (Luck & Vogel, 2013; Luria & Vogel, 2011) and constrained eye movement paradigms (Theeuwes et al., 2009). Future experiments could parametrically vary the demands of the task, as done in hybrid search tasks, in order to explore this hypothesis.

The present study describes the effects of the task in two timescales: first, the local processing of information from each fixation using fRPs and then, the global processing of information throughout the trial, using mass univariate deconvolution modelling to reduce the effects of overlapping brain activity in successive fixations. These results stress the importance of the goal in free-viewing experiments, and the interaction between different timescales in complex sequential tasks as it occurs when sampling almost any scene -static or dynamicin the real world. Future work should explore more deeply this interaction. One possibility would be to parametrically manipulate a relevant cognitive variable, such as asking participants to memorise different numbers of items (Wolfe, 2012). Another alternative would be to manipulate the task’s goal by finishing trials in different ways (e.g. button press). Altogether, concurrent EEG and eye movements recording is an ideal setup to explore the interaction between events and processes in different timescales and hierarchies of a complex task. This is particularly relevant when studying behaviour in natural environments, and is arguably a necessary next step in neurocognitive research.

## Abbreviations

EEG: Electroencephalogram, electroencephalography;
ERP: Event-Related Potential;
FRP: Fixation-Related Potential;
TRF: Temporal Response Function;
VS: Visual Search;
AS: Active Search;
TP: Target Present;
TA: Target Absent;
EX: Exploration;
TPS: Trial Progression Score;
ICA: Independent Component ANalysis;
PCA: Principal Component Analysis;
NTO: non-target objects;
NTF: non-target faces;
TFCE: Threshold-Free Cluster Enhancement method

## Acknowledgements

D.C. was supported by the CONICET, M.J.I was supported by the University of Nottingham, J.E.K. was supported by the CONICET and the University of Buenos Aires (UBA). J.E.K received research grants from CONICET (PIP 11220150100787CO) and ANPCyT (PICT 2018-2699). J.E.K. and M.J.I. received an award from ARL (Cooperative Agreement Number W911NF1920240 and W911NF2120237). We thank Susannah Bilsborough and Joseph Conyers for their collaboration with the data acquisition and Anthony Ries for insightful discussions.

## Code and Data availability statement

The code and sample data are available at *[a LINK that will be available upon publication]*. Raw data will be available upon request.

## Author contributions

M.J.I. and J.E.K. designed and programmed the experiment, collected the data and supervised the study. D.C., M.J.I. and J.E.K. preprocessed the EEG and eye-tracking data. D.C., M.d.F. and J.E.K. performed the deconvolution analysis. D.C., M.d.F., M.J.I and J.E.K. discussed the results and wrote the manuscript. All authors reviewed the manuscript.

## Footnotes

1 https://www.biosemi.com/faq/cms&drl.htm

2 http://pics.stir.ac.uk/2D_face_sets.htm

3 https://kdef.se/home/about%20akdef.html

4 http://vision.ucsd.edu/content/yale-face-database

5 https://www.cl.cam.ac.uk/research/dtg/attarchive/facedatabase.html

6 http://wiki.cnbc.cmu.edu/Face_Place

7 http://pics.stir.ac.uk/2D_face_sets.html

8 http://olivalab.mit.edu/MM/uniqueObjects.html

9 http://github.com/Mensen/ept_TFCE-matlab

