## Supporting material for "Active search signatures in a free-viewing task exploiting concurrent EEG and eye movements recordings"

### Supplementary Methods

#### Single-participant (first level) statistical analysis

As described in the Methods section, we ran mass univariate regressions on the continuous EEG data, using treatment coding. To clarify how the interpretation of the responses depends on the reference level of the categorical effect, we built a synthetic response simulating categorical effects *A*, *B* and *C* and the interaction with a continuous effect *x* (Fig 2). Effect B had a larger positive peak than effect A, while effect C had a larger negative peak than effect A, followed by a square-form continuous interaction at later times. When effect C is coded as 1, the estimation of the base condition response (no increase in the positive and negative peaks and no interaction) is smaller and the parameter C has the expected sign associated with the effect C. On the contrary, switching the reference level for C gives a stronger negative peak in the estimation of the base condition (intercept), since now effect C is added to the base condition, the corresponding parameter has the opposite sign. The parameter $\beta_C$ for the variable $X_C$ now estimates the difference between FRPs of the new base condition and the condition when C effect is absent, which is equivalent to effect A.

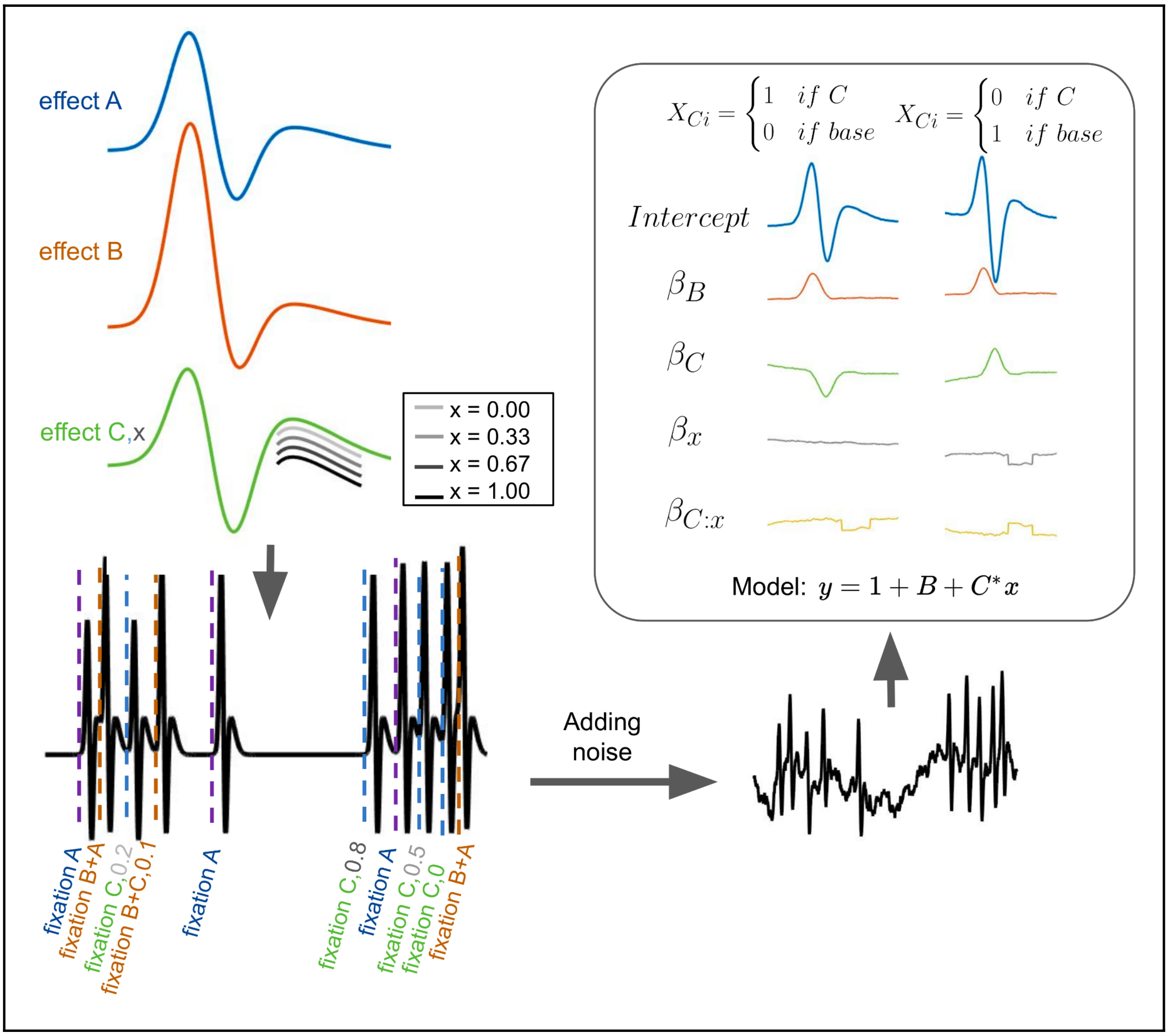

**Supplementary Figure 1. Interpretation of parameters in treatment coding**. Synthetic responses A, B and C, x were set along time simulating the distribution of fixation events for an experimental subject. Pink noise was then added to the signal. The synthetic signal can be deconvolved with a linear model, leading to the beta values shown in the top right box. Notice the differences in values and signs, depending on the chosen reference.

### Models notation

Throughout the manuscript, we adopted the 'symbolic' notation introduced by Wilkinson and Rogers (Wilkinson & Rogers, 1973). This was first proposed in the context of applied statistics to provide a concise and flexible notation for specifying complex (linear and non-linear) models. This notation has become the standard in some fields and applications, being the default notation used by MATLAB Statistics and Machine Learning toolbox, and in statistical models in R (for instance, see (Bates et al., 2014)), which are heavily used in eye movement analysis. Differences in notation often affect the use of models with similar equations in different fields. This is particularly prominent in neuroscience, where many contributions come from diverse fields including engineering, statistics, and computer sciences. In this case, the equations can also be written in linear systems terms (Crosse et al., 2016; Lalor et al., 2009). The equation for the full model (eq. 4) can be presented as:

$$EEG(t) = h_1(t) * s_1(t) + h_{category}(t) * s_{category}(t) + h_{task}(t) * s_{task}(t) + h_{TPS}(t) * s_{TPS}(t) + h_{task:TPS}(t) * s_{TPS}(t) * s_{task}(t) \quad (5)$$

where the *h(t)* functions are the temporal response function (TRFs) to each variable of interest. And the s(t) can be defined as,

$$s_{TPS}(t) = \sum_i \delta(t - t_i) * TPS_i \quad (6)$$

where $TPS_i$ is a continuous value between 0 and 1, and the *δ()* is the Dirichlet function that is 1 when the argument is 0, and 0 for any other value.

$$s_{task}(t) = \sum_i \delta(t - t_i) * task_i \quad (7)$$

where $task_i$ is 0 during EX and 1 during VS, and similarly, the $s_{category}$ is defined category, equal to 0 when it is an object and equal to 1 when it is a face.

$$s_1(t) = \sum_i \delta(t - t_i) \quad (8)$$

The $s_1$ corresponds to the reference level, and it is present every time there is a fixation, even when all the other levels *TPS, task, and category* are set to 0.

<u>References:</u>

Bates, D., Mächler, M., Bolker, B., & Walker, S. (2014). Fitting Linear Mixed-Effects Models using lme4. In *arXiv [stat.CO]*. arXiv. http://arxiv.org/abs/1406.5823

Crosse, M. J., Di Liberto, G. M., Bednar, A., & Lalor, E. C. (2016). The Multivariate Temporal Response Function (mTRF) Toolbox: A MATLAB Toolbox for Relating Neural Signals to Continuous Stimuli. *Frontiers in Human Neuroscience*, *10*, 604. https://doi.org/10.3389/fnhum.2016.00604

Lalor, E. C., Power, A. J., Reilly, R. B., & Foxe, J. J. (2009). Resolving precise temporal processing properties of the auditory system using continuous stimuli. *Journal of Neurophysiology*, *102*(1), 349–359. https://doi.org/10.1152/jn.90896.2008

Wilkinson, G. N., & Rogers, C. E. (1973). Symbolic description of factorial models for analysis of variance. *Journal of the Royal Statistical Society. Series C, Applied Statistics*, *22*(3), 392. https://doi.org/10.2307/2346786

### Supplementary Figures

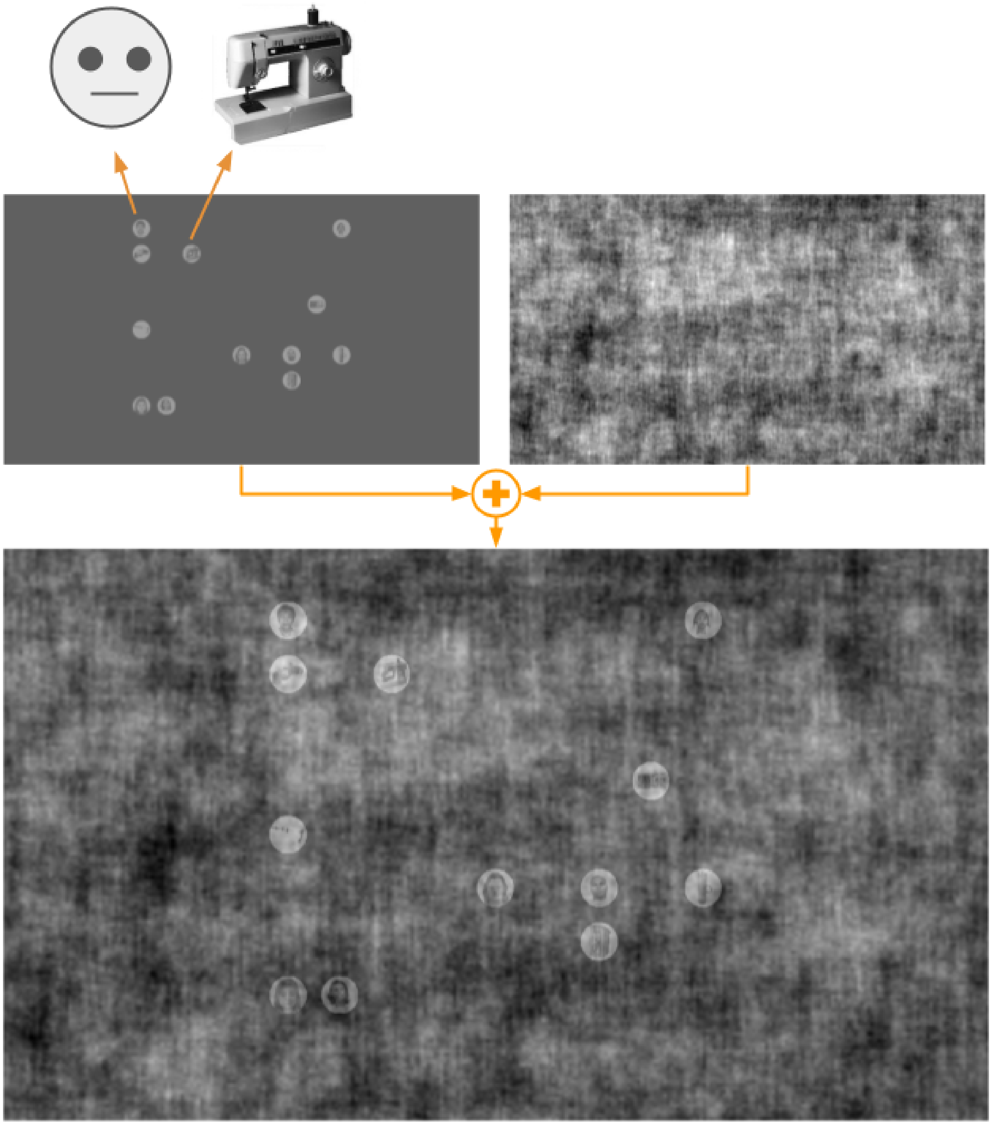

**Supplementary Figure 2. Stimulus example.**.The individual stimuli were randomly placed in a grid, and embedded in a noisy background (shuffled natural scenes). The individual stimuli and the background were made isoluminant. THIS FIGURE WAS MODIFIED FOR THE BIORXIV VERSION, IN THE ORIGINAL EXPERIMENT THE SMILEY FACE IS REPLACED BY A HUMAN FACE.

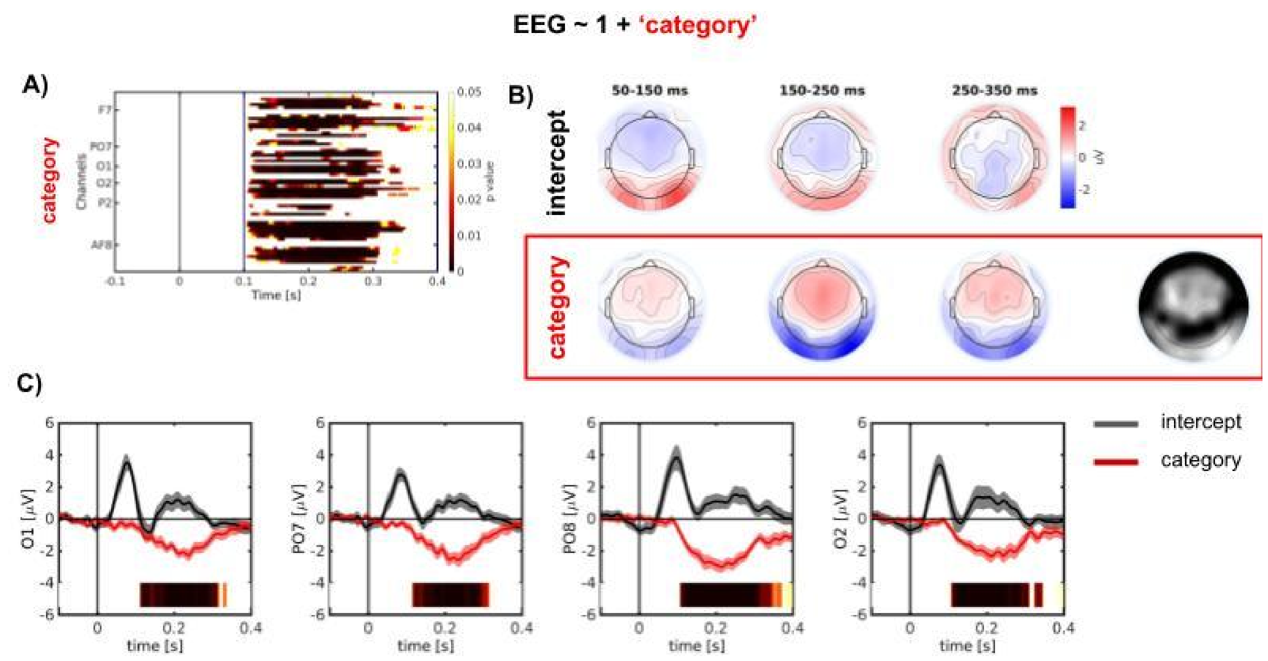


**Supplementary Figure 3. Time-course of category effect on FRPs for visual search (pre and post-target fixations) and exploration tasks (model Eq. 1). A.** Significant samples obtained with the TFCE test. Vertical lines represent the start and end of the significant cluster (with p<0.01). **B.** Scalp topography for different stages of the intercept and the category effect (beta values). The right scalp topography shows the proportion of significant samples per channel during the interval of the significant cluster (vertical bars in A). The colour scale represents activation from 0 (black) to the whole interval (white). **C.** Mean and s.e.m. across participants of the intercept (black) and the category (red) effect for selected channels [O1, PO8, PO7, O2]. The colour bar below each panel represents the significant intervals after TFCE second-order statistics applied to the face-object (category) effect. The intensity of the bar represents the significance level (p-value), only for the p-values < 0.05, the scale is the same as in panel A. The shades represent the standard error of the mean across participants. Number of fixations = 8654. Number of participants = 16.

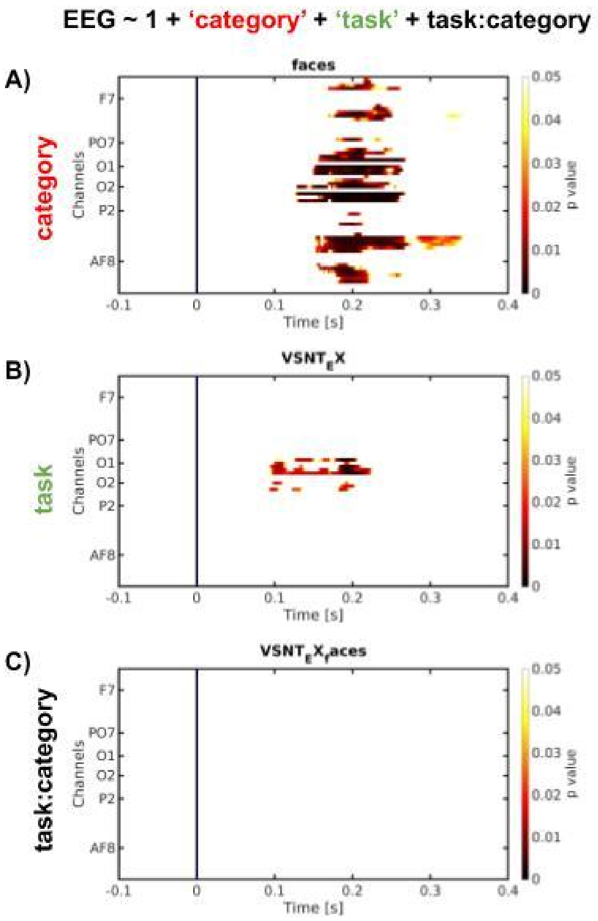


**Supplementary Figure 4. Time-course of category and task effects and their interaction on FRPs for active search and exploration tasks (*EEG ~ 1 + category + task + task:category*).** Significant samples obtained with the TFCE. Number of fixations = 5578. Number of participants = 16.

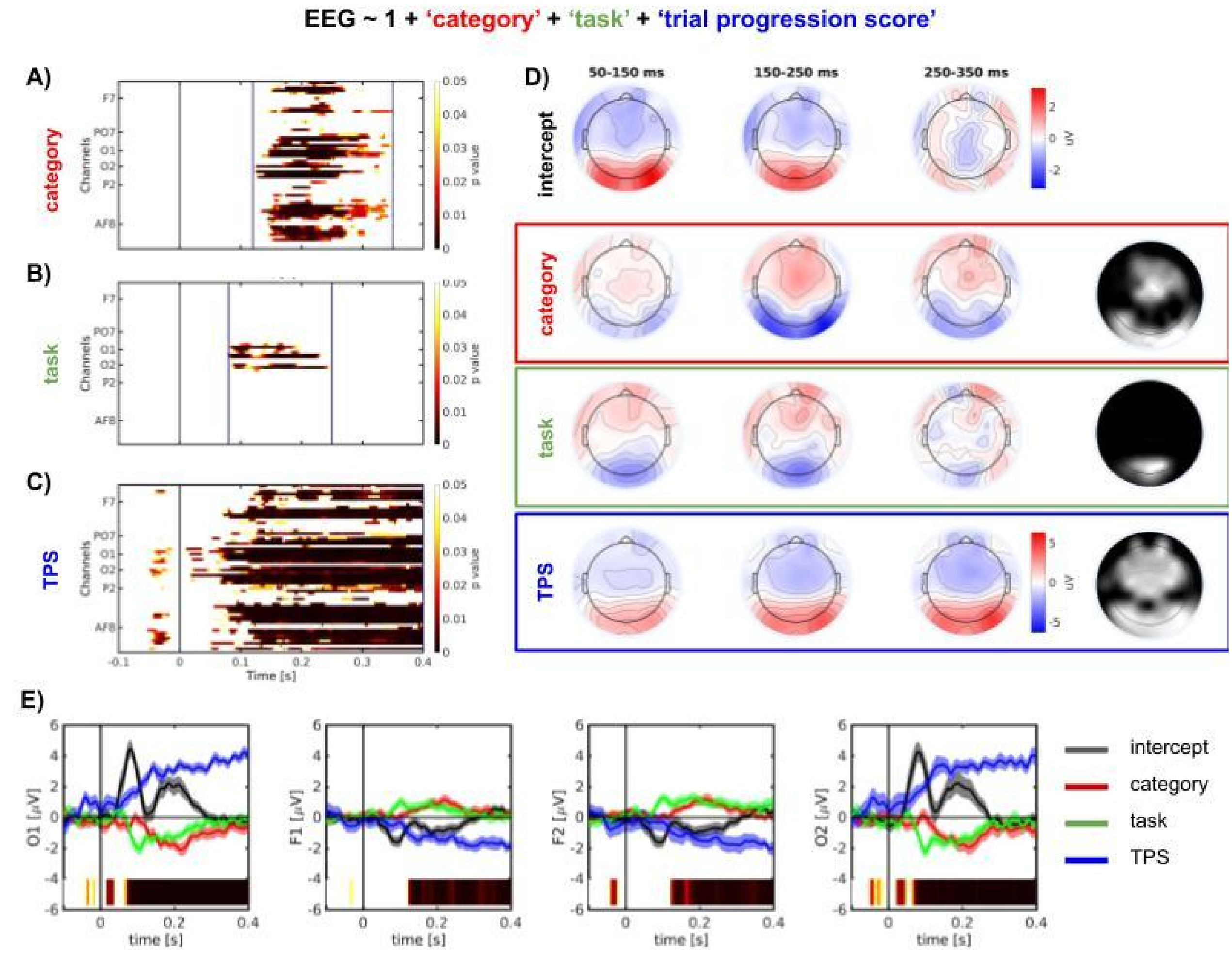


**Supplementary Figure 5. Time-course of trial progression score TPS) effect on FRPs for active search and exploration tasks (model Eq. 3). A.B.C.D.** Significant samples obtained with the TFCE test for the category (**A**), the task (**B**) and the TPS (**C**) effect. Vertical lines represent the start and end of the significant cluster (with $p<0.01$). **D.** Scalp topography for different stages of the intercept and the category effect (beta values). The right scalp topography shows the proportion of significant samples per channel during the interval of the significant cluster (vertical bars in A-C). The colour scale represents activation from 0 (black) to the whole interval (white). **E.** Mean and s.e.m. across participants of the intercept (black), the category (red), the task (green) and the TPS (blue) effect for selected channels [O1, F1, F2, O2]. The colour bar below each panel represents the significant intervals after TFCE second-order statistics applied to the TPS effect. The intensity of the bar represents the significance level (p-value), only for the p-values $< 0.05$, the scale is the same as in panel A. The shades represent the standard error of the mean across participants. Number of fixations = 5578. Number of participants = 16.

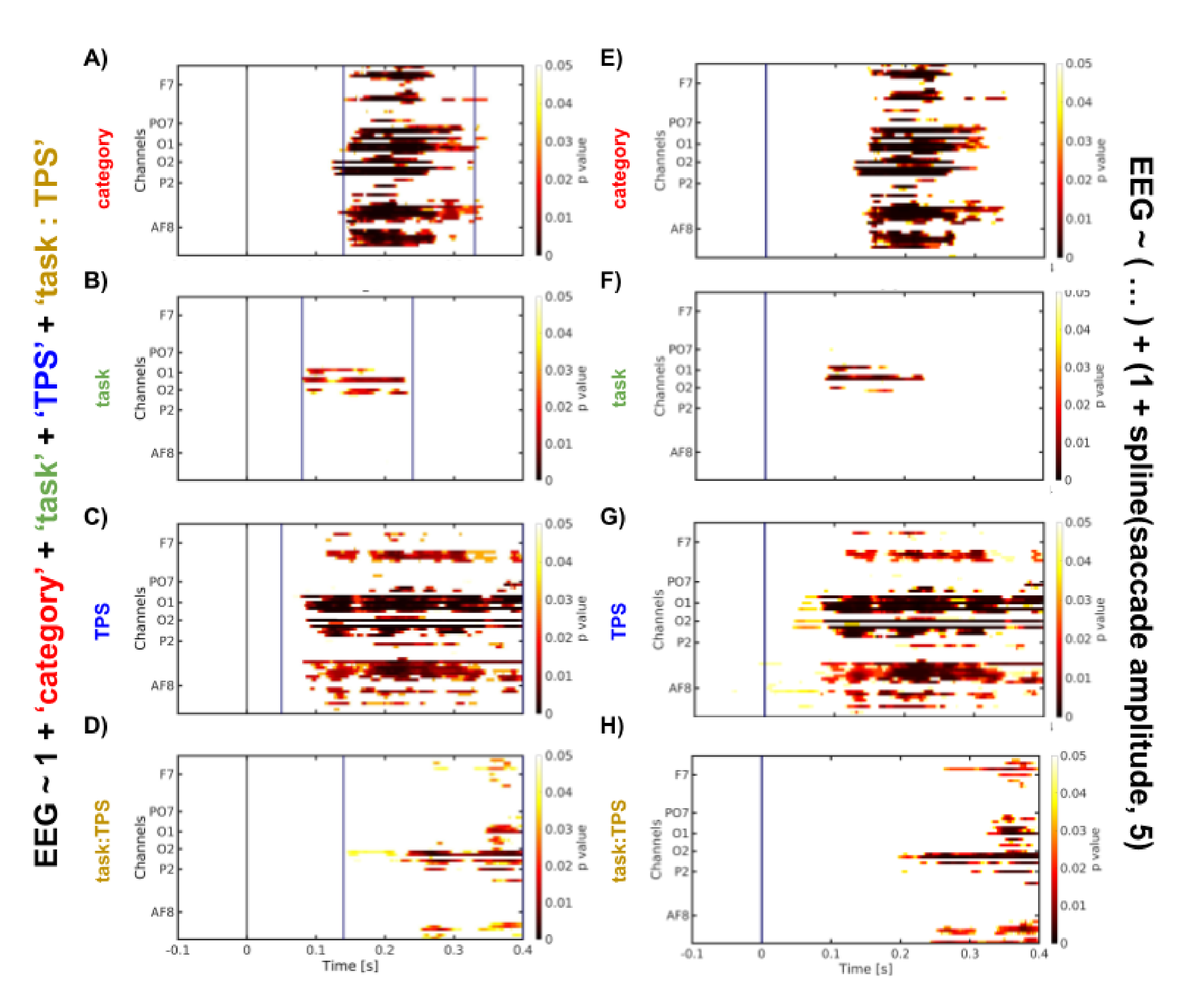


**Supplementary Figure 6. Expanded model with saccade events as variables. A.B.C.D.** Time-course of trial progression score (TPS) and the interaction with the task effect on FRPs for active search and exploration tasks (model Eq. 4; same as Fig. 5). Significant samples obtained with the TFCE test for the category (**A**), the task (**B**), the TPS effect (**C**), the interaction between TPS and the task (**D**). **E.F.G.H.** Same as A.B.C.D. but with the addition of an intercept and a fifth-order spline on the saccade amplitude aligned to the onset of the saccades previous to every valid fixation: *fixation related activity ~ ( 1 + category + task + TPS + task:TPS )* ; and *saccade related activity ~ ( 1 + spl(saccade amplitude, 5) )*. Number of fixations = 5578. Number of participants = 16.

### Graphical Abstract

We concurrently recorded EEG and eye movements to study the brain responses to different categories (faces and objects), during different tasks (visual search and exploration). By applying a deconvolution analysis approach, we were able to estimate the contribution of the different elements embedded in the task. These results broaden our understanding of visual information processing in free viewing and contribute towards understanding the brain under closer to real-life conditions.

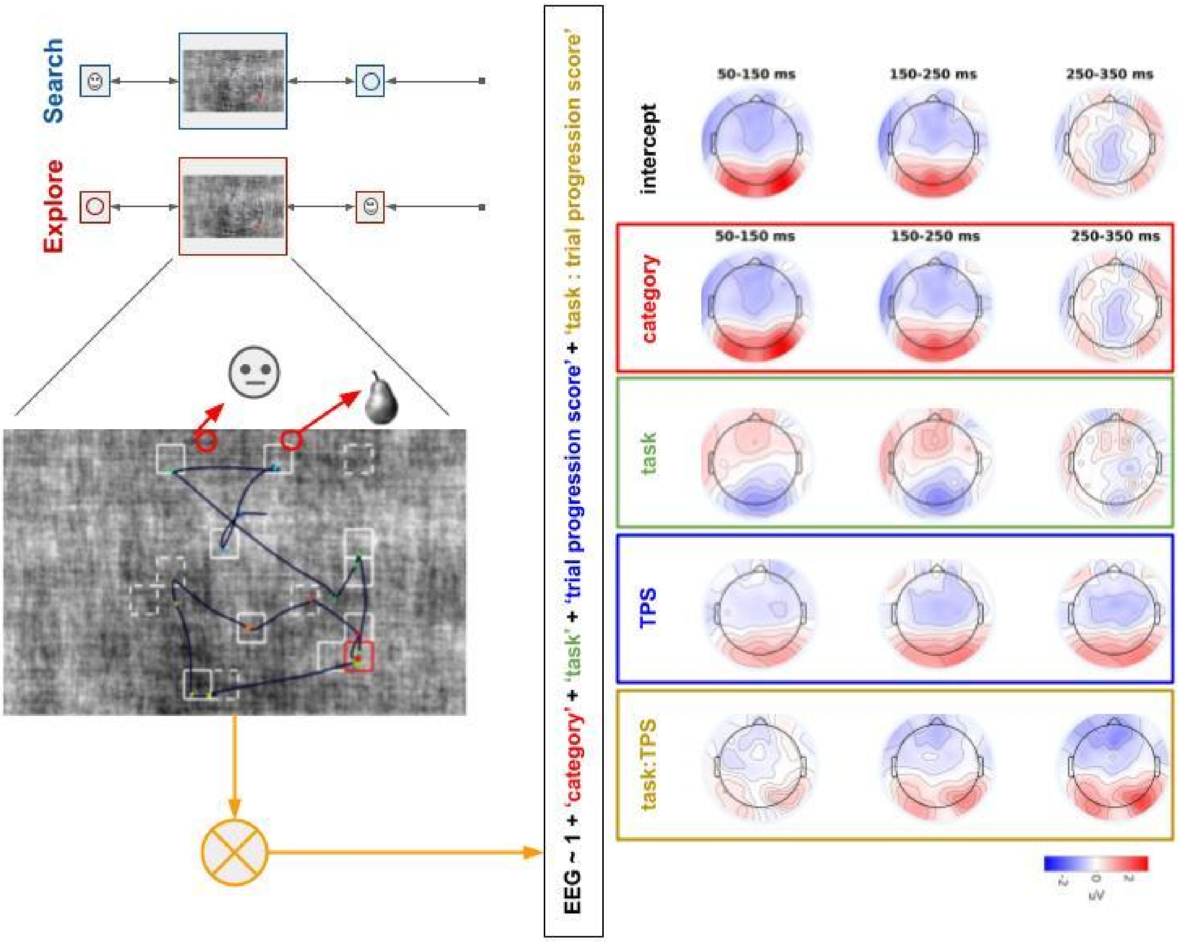
